# Epigenetic Conditioning Induces Intergenerational Resilience to Dementia in a Mouse Model of Vascular Cognitive Impairment

**DOI:** 10.1101/2021.06.16.448715

**Authors:** Krystal Courtney D. Belmonte, Eleanor B. Holmgren, Tiffany A. Wills, Jeff M. Gidday

## Abstract

**Background:** Vascular cognitive impairment and dementia (VCID), which occurs immediately or in delayed fashion in 25-30% of stroke survivors, or secondary to chronic cerebral hypoperfusion, is the second leading cause of dementia following Alzheimer’s disease. To date, efficacious therapies are unavailable. We have shown previously in mice that repetitive hypoxic preconditioning (RHC) induces a long-lasting resilience to acute stroke (Stowe et al., 2011). More recently, we documented that untreated, first-generation adult progeny of mice exposed to RHC prior to mating are protected from retinal ischemic injury (Harman et al., 2020), consistent with accumulating evidence supporting the concept that long-lasting phenotypes induced epigenetically by intermittent stressors may be heritable. We undertook the present study to test the hypothesis that RHC will induce resilience to VCID, and that such RHC-induced resilience can also be inherited.

**Methods:** Chronic cerebral hypoperfusion (CCH) was induced in C57BL/6J mice secondary to bilateral carotid artery stenosis with microcoils in both the parental (F0) generation, and in their untreated first-generation (F1) offspring. Cohorts of F0 mice were directly exposed to either 8 wks of RHC (1 h of systemic hypoxia [11% oxygen] 3x/week) or normoxia prior to CCH. Cohorts of F1 mice were derived from F0 mice treated with RHC prior to mating, and untreated, normoxic controls that were age-matched at the time of stenosis induction. Demyelination in the corpus callosum of F0 mice was assessed following 3 months of CCH by immunohistochemistry. Mice from both generations were assessed for short-term recognition memory *in vivo* by novel object preference (NOP) testing following 3 months of CCH, and a month thereafter, *ex vivo* measurements of CA1 hippocampal long-term potentiation (LTP) were recorded from the same animals as a metric of long-lasting changes in synaptic plasticity.

**Results:** Three months of CCH caused demyelination and concomitant impairments in recognition memory in control mice from both generations. However, these CCH-induced memory impairments were prevented in F0 animals directly treated with RHC, as well as in their untreated adult F1 progeny. Similarly, hippocampal LTP was preserved in the 4-month CCH cohorts of mice directly treated with RHC, and in their untreated offspring with CCH.

**Conclusions:** Our findings demonstrate that RHC or other repetitively-presented, epigenetic-based therapeutics may hold promise for inducing a long-lasting resilience to VCID in treated individuals, and in their first-generation adult progeny.

## INTRODUCTION

Vascular contributions to cognitive impairment and dementia (VCID), secondary to chronic cerebral hypoperfusion (CCH), acute stroke, and other causes, is the second most common form of dementia after Alzheimer’s disease (AD) (O’Brien & Thomas, 2015; Gorelick et al., 2016b; Kalaria et al., 2016; Iadecola et al., 2019). Moreover, the prevalence of cerebrovascular disease is in fact higher than the prevalence of AD (Heron, 2019), further increasing the probability that many cases of VCID remain undocumented. To date, there is no efficacious therapy for VCID.

An epigenetics-based VCID therapeutic strategy may be able to meet this clinical challenge. Specifically, preconditioning the brain to activate changes in gene expression is a well-established approach to promote endogenous pathways of cell survival and tissue resistance to injury (Sprick et al., 2019; Hao et al., 2020). We and others have previously shown the efficacy of hypoxic preconditioning in inducing protection against acute stroke (Miller et al., 2001; Stowe et al., 2011; Wacker et al., 2012; Gidday, 2015). Importantly, our lab has also documented, in a mouse model of focal stroke, that the duration of this inducible neurovascular-protective phenotype can be extended dramatically - from days to months - following the last of a series of repetitive hypoxic conditioning (RHC) stimuli (Stowe et al., 2011). In a recent study, we showed that 4 months of RHC in male and female mice, prior to breeding, results in a heritable resilience to retinal ischemic injury in their untreated, first-generation progeny (Harman et al., 2020).

While there is considerable precedent in the literature for so called ‘intergenerational epigenetic inheritance’ in animals, and in humans as well (Pembrey et al., 2014; Bowers & Yehuda, 2016), by far, the vast majority of this research is focused on demonstrating the heritability of disease risk and disease itself. In the present study, we hypothesized that RHC will induce resilience to cognitive deficits in a mouse model of VCID, and that this dementia-resilient phenotype will be inherited. The data we collected support our hypothesis, documenting the therapeutic potential of activating an endogenous resiliency against dementia capable of promoting healthier lives that can span generations.

## METHODS

### Animals

All animal procedures were approved by our institutional Animal Care and Use Committee and performed in accordance with the ARRIVE Guidelines and the NIH Guide for the Care and Use of Laboratory Animals. Male and female C57BL/6J mice were purchased from The Jackson Laboratory (Bar Harbor, ME, USA) at 8-9 wks of age. Animals were allowed to habituate to the animal facility for 1 wk prior to treatment, kept on a 12 h/12 h light/dark cycle with food and water provided ad libitum, and randomly assigned into experimental groups following ~2 wks of habituation.

### Experimental Design

Mice were treated with 8 wks of RHC, in parallel with age-matched normoxic controls. Separate cohorts were used for F0-generation studies of chronic cerebral hypoperfusion (CCH), and for F1-generation studies, by breeding within their respective RHC or normoxic groups to produce F1-generation progeny. Specifically, males treated with 8 wks of RHC were paired with 8-wk RHC-treated females to produce the [F1: RHC]* generation (asterisk denoting that F1-generation mice were never exposed to RHC themselves), and normoxic males and females were paired to produce the F1: CTL generation (Figure 1). In all cases, both F0 parents remained in the cage until their F1 offspring were weaned at PND21, and housed thereafter in standard cages, breathed ambient air throughout their lives, and were age-matched with their respective F0 generation at the time of carotid microcoil placement, neurocognitive testing, and the collection of hippocampal slices for LTP measurement.

**Figure 1.**
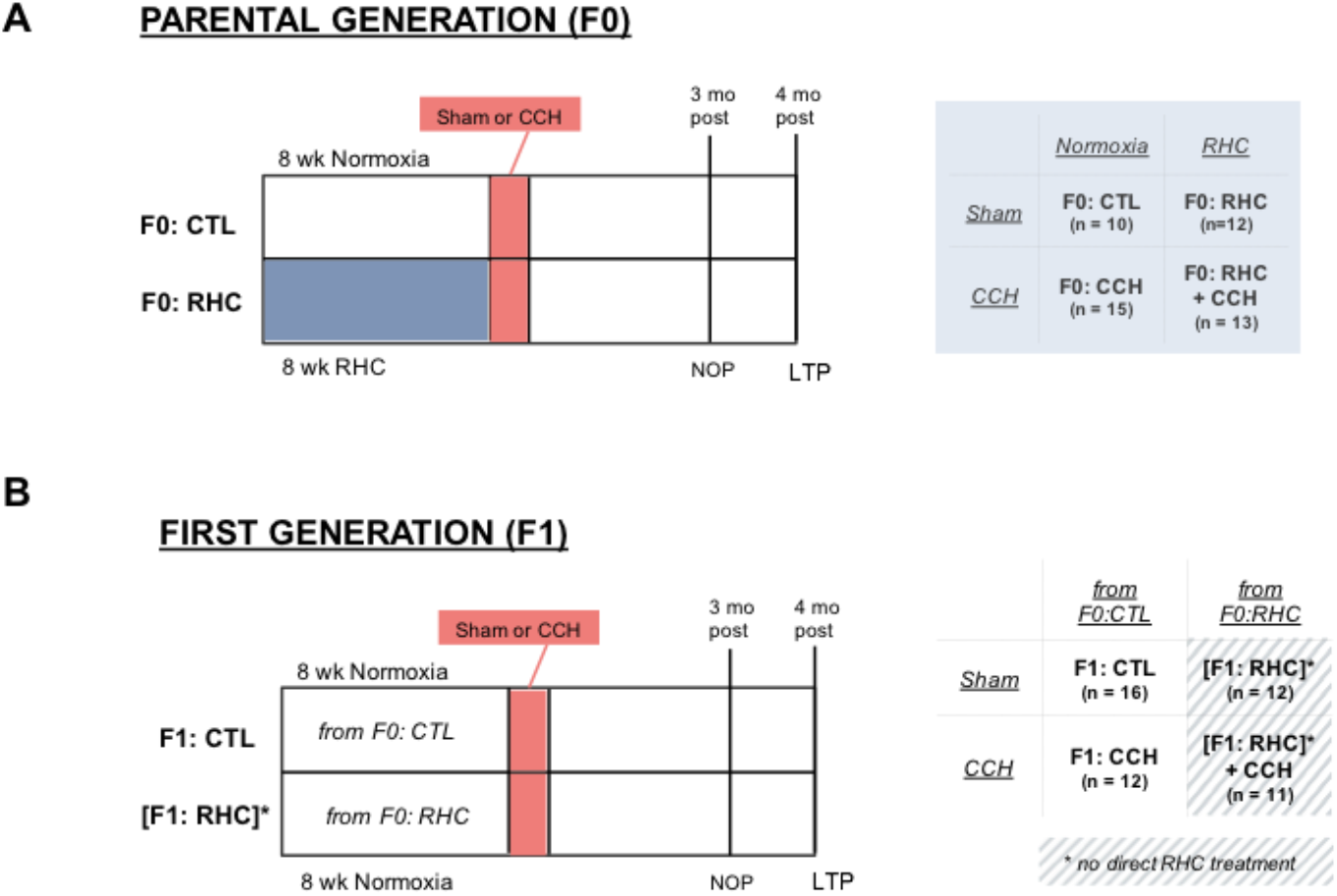
Experimental design for repetitive hypoxic conditioning (RHC) studies in mice with chronic cerebral hypoperfusion (CCH). (A) Parental generation male and female adult mice were exposed to repetitive hypoxic conditioning (11% O_2_ for 1 h every other day), or normoxia, for 8 wks. After the last treatment, mice were either subjected to bilateral carotid artery stenosis (BCAS) surgery to induce chronic cerebral hypoperfusion (CCH), or sham surgery. (B) First generation offspring were derived from a separate cohort of F0 animals, both males and females, that were only subjected to either 8 wks of RHC, or normoxia, prior to mating. Male and female F1 offspring originating from these pairings never received RHC directly. At ~20 wks of age, all F1-generation animals were subjected to either CCH or sham surgery. Novel object preference (NOP), a neurobehavioral measurement of object recognition memory, was performed after 3 mo of CCH, and hippocampal long-term potentiation (LTP) was assessed as an electrophysiological proxy of memory-associated synaptic plasticity, approximately 1 mo later.

### Repetitive Hypoxic Conditioning

Repetitive hypoxic conditioning (RHC) involved exposing mice to mild-to-moderate systemic hypoxia (11% O_2_) using an oxygen tension-controlled chamber (Biospherix, Parish, NY, USA). The animal’s home cages were placed directly in the chamber, abrogating a change in environment other than ambient air, and were allowed free access to food and water during treatment. Mice were treated for 1 h, every other day, for 8 continuous wks. Age- and sex-matched mice exposed to normal atmospheric oxygen served as normoxic controls. In both studies, only the F0 generation was exposed to RHC.

### Chronic Cerebral Hypoperfusion

To model the effects of vascular cognitive impairment and dementia, chronic cerebral hypoperfusion (CCH) was achieved by bilateral carotid artery stenosis (BCAS) in accordance with prior publications (Shibata 2004, Shibata 2007, Nishio 2010). In brief, at ~20 wks of age, control and experimental animals were anesthetized with ketamine (100 mg/kg) and xylazine (10 mg/kg) intraperitoneally, and injected with sustained-release buprenorphine (0.1 mg/kg) prior to surgery. Through a midline cervical incision, both common carotid arteries were exposed, lifted using 2-0 surgical thread, and isolated from their vagus nerves. Gold-plated microcoils (0.18-mm internal diameter; 0.50 mm pitch; 2.5-mm total length; Motion Dynamics Corporation, Fruitport, MI, USA) were wrapped around both arteries. Forebrain cerebral blood flow (CBF) was recorded from the carotid artery bilaterally, prior to the application of the first microcoil and following application of the second microcoil, by laser Doppler flowmetry (BLF21; Transonic Systems Incorporated, Ithaca, NY, USA). Sham animals were exposed to the same anesthesia and operative procedures as CCH mice, but without microcoil placement. Reductions in CBF resulting from coil placement were normalized to coil pre-application CBF values for each carotid. Incisions were closed using Vetbond tissue adhesive (3M, Saint Paul, MN, USA). Body temperature was maintained during the operative procedure and post-operative recovery with a warming pad. Animals were placed in a new home cage post-surgery and closely monitored till ambulatory. Animals were single-housed to avoid any rupturing of the incision closures by other littermates, and were provided with IACUC-approved supplementary nesting enrichment required for single-housing purposes. Animals were then left undisturbed in our vivarium for 3 months until neurocognitive testing was initiated.

### Demyelination

Given previous reports documenting a loss of myelin integrity in mice after 1 mo of CCH (Shibata et al., 2004; Shibata et al., 2007; Miki et al., 2009; Coltman et al., 2011; Ahn et al., 2016; Ben-Ari et al., 2019), we assessed myelin density in the corpus callosum after 3 months of CCH in F0 cohorts of mice with and without RHC treatment, using a well-established fluorescent histochemical method (Kanaan 2006; Ahn 2016). In brief, brains were removed after 4 mo of CCH (or sham surgery) and post-fixed in Z-Fix (Anatech Limited, Battle Creek, MI, USA) for 24 h at 4°C before cryoprotection in 30% sucrose. Prior to sectioning, brains were flash-frozen with isopentane at −20°C. Serial, 20μm-thick coronal sections were obtained from the anterior brain (approximately 0.86 mm posterior to Bregma, per mouse brain atlas) using a cryostat at −11°C, and stored at 4°C in PBS. Free-floating sections were rinsed in 0.2% Triton X-100 PBS (PBSt) for at least 20 min at room temperature before staining with a fluorescent myelin stain (FluoroMyelinTM Green; 1:300 in PBS; Molecular probes, Eugene, OR, USA) for 20 min at room temperature. Seven to eight sections per animal were mounted onto SuperFrost glass slides (Thermo-Fisher Scientific; Walthan, MA, USA) with mounting medium (ProLong Gold anti-fade; Invitrogen, Carlsbad, CA, USA), cover-slipped, and visualized using an Olympus BX51 fluorescent microscope (Olympus Life Science, Waltham, MA, USA). Images were captured at 4x magnification using cellSense imaging software (Olympus Life Science, Walthan, MA, USA). The entire corpus callosum was selected (Figure 2A) using the “freehand selection” option and pixel intensity for the 7-8 sections was determined bilaterally and averaged using ImageJ 2 software (NIH). Sections from representative animals within each experimental group were always stained concomitantly, which permitted intensities for each experimental group to be normalized to sham controls.

**Figure 2.**
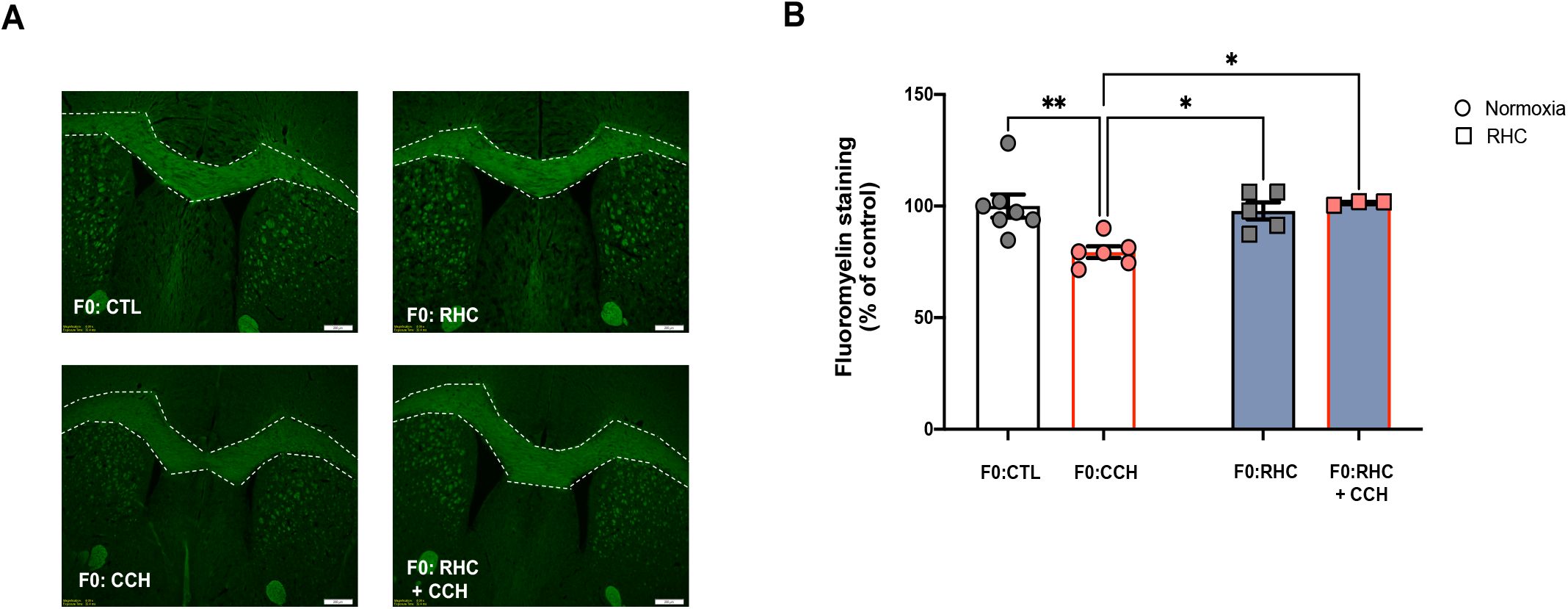
Repetitive hypoxic conditioning (RHC) reduces white matter injury in the corpus callosum following 3 mo of chronic cerebral hypoperfusion (CCH). (A) Representative fluoromyelin images in coronal brain sections from F0 mice following 3 mo of CCH or sham surgery (CTL). Region of corpus callosum used for quantification of staining intensity is delineated by white lines. (B) Fluoromyelin staining intensity, used as a metric for corpus callosum myelination, was normalized for each experimental group to untreated controls (F0: CTL). *p<0.05, **p<0.01, by two-way ANOVA with Tukey’s post-hoc analysis (n=3-7/group). Mean±SEM. Scale bar 200 μm.

### Novel Object Preference

Novel object preference (NOP) testing is a well-established neurobehavioral tool to assess recognition memory (Antunes & Biala 2012; Vogel-Ciernia et al., 2014; Lueptow et al., 2017; Morena-Castilla et al., 2018). In the VCID model we used, impairments in recognition memory have been confirmed by NOP following 1 month (Patel et al., 2017; Dominguez et al., 2018) of CCH. We commenced NOP testing 3 mo after CCH or sham surgery in both F0 and F1 mice. Mice were individually habituated to a dimly-lit black Plexiglass glass box with a white colored floor (20 x 20 x 20 cm) for 15 min on the day before the test (Figure 3A). The following day, a testing session consisting of two trials was administered - the familiar object exposure, and the novel object exposure. For the former, mice were individually placed in the box containing two identical objects for 10 min and then returned to their home cage. Three hours later, mice were returned to the box, which now contained a novel object along with one of the familiar objects, for the novel object preference assessment (Figure 3A). For 10 min, mice were freely allowed to explore the objects. Time (seconds) spent with each object was recorded using a video camera mounted above the box. Exploration was defined as sniffing or touching either object at a distance of <2 cm from the object. To be counted, the mouse had to be on all 4 feet during the exploration; times during which the mouse sat on or climbed the object were not included. All objects were randomized and counterbalanced across animals. Between each session, the open field box and objects were thoroughly cleaned with 70% ethanol. Analysis of NOP behavior was conducted by video playback by an observer blinded to experimental treatment conditions. Recognition memory was quantified by a discrimination index (DI), as follows: (time exploring novel object - time exploring familiar object) / (time exploring novel + familiar). Thus, a larger discrimination index is indicative of more time exploring the novel object, reflecting the animal’s recognition memory of the familiar object.

**Figure 3.**
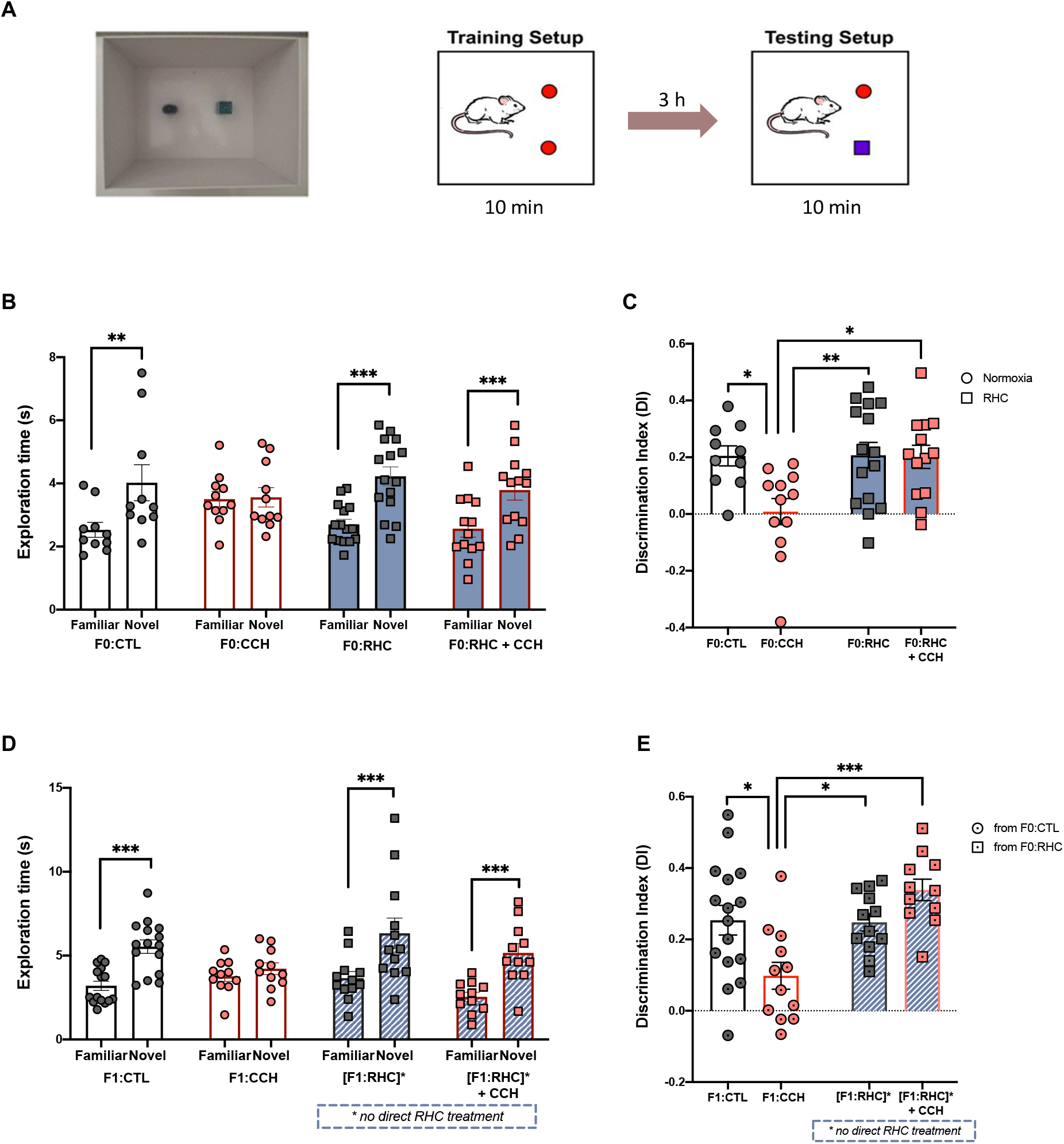
Repetitive hypoxic conditioning (RHC) prevents loss of object recognition memory *in vivo* in F0 mice and their untreated F1 progeny. Schematic diagram of the spontaneous object recognition testing paradigm (A). Familiar and novel object exploration times and (B) discrimination indices of the four F0 experimental groups (C). Note, impairments in recognition memory following 3 mo of CH were prevented in F0 mice directly treated with RHC. Familiar and novel object exploration times (D) and discrimination indices (E) of the four F1 experimental groups. Note, in parallel with findings in F0 mice (B,C), loss of recognition memory following 3 mo of CCH is prevented in untreated F1 mice derived from F0 parents treated with RHC prior to mating (D,E). *p<0.05, **p<0.01, ***p<0.001, by paired, non-parametric Wilcoxon test (B,D) and two-way ANOVA with Tukey’s post-hoc analysis (C,E; n=10-16/group). Mean±SEM.

### Hippocampal Long-term Potentiation

Long-term potentiation (LTP) refers to a strengthening of synaptic connectivity following co-activation of neurons in a network, and is a well-established form of synaptic plasticity that subserves memory function in the mammalian hippocampus, amygdala, and other cortical brain structures (Nicoll 2017; Sumner et al., 2020). Hippocampal slices were prepared from mice after 4 mo of CCH (or sham surgery), a month following the completion of NOP testing for that animal. Mice were anesthetized with isoflurane and decapitated. Brains were quickly removed from the skull, and 300-μm thick hippocampal slices were prepared using a Tissue Slicer (Leica Biosystems Inc., Buffalo Grove, IL), collected in artificial cerebrospinal fluid (ACSF) (125 mM NaCl, 4.4 mM KCl, 2 mM CaCl_2_, 1.2 mM MgSO_4_, 1 mM NaH_2_PO_5_, 10.0 mM glucose and 26.0 mM NaHCO_3_; saturated with 95% O_2_ and 5% CO_2_), and transferred to a submerged recording chamber where they were perfused with heated (28°C) and oxygenated (95% O_2_ - 5% CO_2_) ACSF. Slices were allowed to equilibrate at ACSF for at least 1 h before recordings were initiated. A bipolar stainless steel stimulating electrode and a borosilicate glass recording electrode filled with ACSF were both positioned in the stratum radiatum of the hippocampal CA1 region to elicit and record extracellular field responses (Figure 4A). The field excitatory postsynaptic potential (fEPSP) slope was monitored using Axon Patch-Clamp instrumentation (Molecular Devices, San Jose, CA, USA), and recordings were analyzed with Clampfit software (Molecular Devices, San Jose, CA, USA). The test stimulation strength was determined for each input as the current needed to elicit a field EPSP of 40% maximal slope. Baseline fEPSPs were recorded for 20 min. LTP was induced with a strong tetanization protocol consisting of two stimulus trains (100 biphasic constant-current pulses per train at 100 Hz, inter-train interval 20 sec). Field excitatory postsynaptic potentials were recorded for 60 min post-tetanus and results displayed as the averaged fEPSPs recorded over the last 20 min post-tetanus.

**Figure 4.**
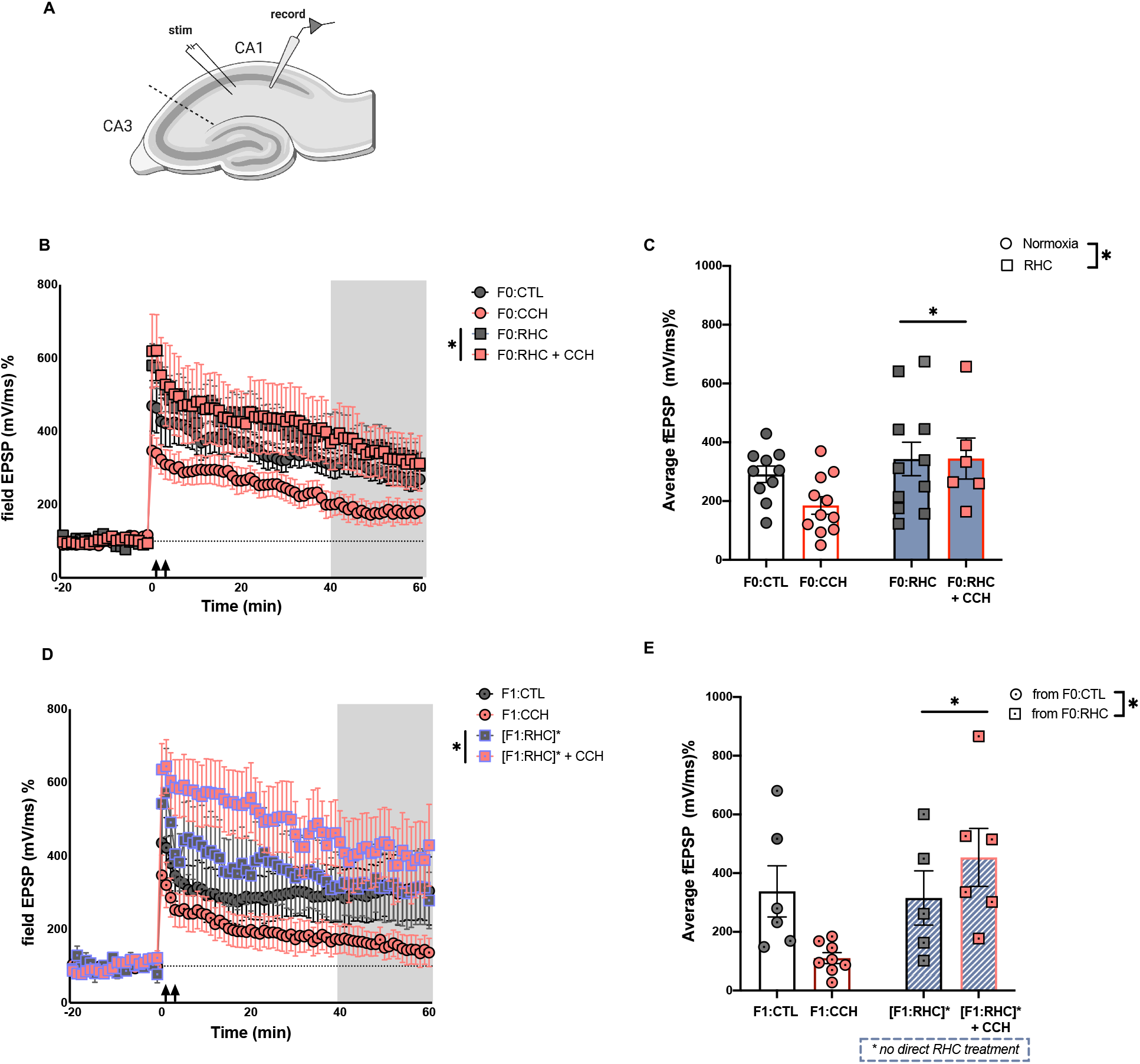
Repetitive hypoxic conditioning (RHC) preserves *ex vivo* long-term potentiation (LTP), a metric of synaptic plasticity, in CA1 hippocampus of F0 mice and their untreated F1 progeny. (A) Schematic of a coronal hippocampal slice showing positions of the stimulating (stim) electrode and recording (record) electrode. (B) Field excitatory postsynaptic potential recordings (fEPSPs), normalized to their respective baselines, for the 4 experimental groups of F0 mice; the increase in fEPSP following tetanus stimulation (double arrows) indicates a hippocampal LTP response. The average fEPSP over 40-60 min post tetanus (shaded in grey) of each F0 group is represented in (C). Similarly, fEPSP responses (D), and their average amplitude over 40-60 min (E), are shown for the 4 groups of F1 mice. Note, LTP was preserved following 4 months of CCH in F0 mice directly treated with RHC, and in their untreated F1 progeny. *p<0.05 main effect of RHC by two-way ANOVA with Tukey’s post-hoc analysis (n=10-16/group). Mean±SEM.

### Statistical Analyses

GraphPad Prism 9 software (La Jolla, CA, USA) was used to analyze data. For changes in CBF, a two-tailed unpaired t-test was used. White matter myelin densities, discrimination indices for NOP, and field excitatory post-synaptic potential (fEPSP) means from LTP recordings were analyzed by two-way ANOVA with Tukey post-hoc analysis where the factors were treatment (CTL vs. RHC) and injury (Sham vs. CCH). All values are presented as means ± SEM, with p<0.05 considered significant.

## RESULTS

### Repetitive hypoxic conditioning (RHC) did not affect the magnitude of chronic cerebral hypoperfusion (CCH)

To confirm that the extent of cerebral blood flow (CBF) reduction following bilateral carotid artery stenosis (BCAS) was not affected by our RHC therapy, we measured CBF before and after microcoil placement. No statistical differences in baseline CBF values were found between all groups (data not shown), including RHC-treated mice. Likewise, F1 animals derived from RHC-treated mice exhibited no significant differences in baseline CBF compared to their same-generation controls derived from untreated F0 parents. The lack of significant differences between all experimental groups at baseline suggests neither direct nor parental RHC caused any changes that affected resting levels of forebrain blood flow.

Furthermore, whether untreated or with prior RHC treatment, mice exhibited similar reductions in CBF upon microcoil placement. In both generations, there was a significant reduction in CBF (relative to respective baseline values) after the application of the first (−46±3%) and second (−43±4%) coil compared to sham animals (p<0.0001). Moreover, compared to their generational controls, the extent of CBF reduction in RHC-treated animals (−47±2%, −48±2%) and offspring derived from RHC-treated animals (−42±3%, −40±4%) was similar after the application of the first and second microcoil, respectively (p < 0.0001). These results demonstrate that the magnitude of forebrain CBF reduction caused by microcoil placement was similar in all groups, including mice with prior RHC, indicating that any observed protection afforded by RHC was not the result, at least initially, of a less severe perfusion deficit.

### Repetitive hypoxic conditioning (RHC) prevents chronic cerebral hypoperfusion (CCH)-induced white matter injury

As a metric for white matter (WM) injury and protection, we assessed myelin density in the corpus callosum of F0 mice treated with and without RHC after 3 mo of CCH. Two-way ANOVA of fluoromyelin staining intensities revealed a significant RHC x CCH interaction (F_1,17_ = 7.315, p = 0.0150; Figure 2B). Tukey post-hoc analyses confirmed that, compared to control animals, fluoromyelin intensities were significantly reduced following 4 mo of CCH (F0: CTL vs F0: CCH, p = 0.0070), reflecting CCH-induced demyelination. However, animals treated with RHC prior to CCH exhibited significantly greater fluoromyelin intensities compared to CCH animals (F0: RHC + CCH vs F0: CCH, p = 0.0247), and in fact were indistinguishable from control animals (F0: CTL vs F0: RHC+CCH, p = 0.9967), suggesting that RHC completely prevented CCH-induced demyelination.

### Intergenerational prevention of chronic cerebral hypoperfusion (CCH)-induced impairment in object recognition memory by repetitive hypoxic conditioning (RHC)

For each experimental group, differences between the exploration time of the novel object and familiar object was analyzed using the paired, non-parametric Wilcoxon test. Control animals spent significantly more time exploring the novel object compared to the familiar object (F0: CTL familiar vs novel, p = 0.0039; F1: CTL familiar vs novel, p = 0.0001; Figures 3B, 3D). Mice with 3 mo of CCH exhibited no significant differences in exploration time between the novel and familiar object (F0: CCH familiar vs novel, p = 0.5625; F1: CCH familiar vs novel, p = 0.0674; Figures 3B, 3D). In contrast, F0 animals directly treated with RHC, as well as their untreated F1 offspring, spent significantly more time exploring the novel object compared to the familiar object (F0: RHC familiar vs novel, p = 0.0009, [F1: RHC]* familiar vs novel, p = 0.0005), indicating that RHC prevented CCH-induced losses in recognition memory both in animals receiving the treatment, as well as in their untreated offspring (Figures 3B, 3D).

To account for the differences in total exploration time and standardize such total time amongst all groups for comparison, the discrimination index (DI) was used to quantify the animal’s preference between the novel and familiar object. Two-way ANOVA of DI revealed a significant RHC x CCH interaction in the F0 (F_1,46_ = 4.723, p = 0.0349; Figure 3C) and F1 generation (F_1,47_ =11.47, p = 0.0014; Figure 3E). In both generations, Tukey post-hoc analyses confirmed lower discrimination indices (DI) in CCH animals compared to that of their generational controls, indicating an impairment in recognition memory in the former (F0:CTL vs F0:CCH, p = 0.0228; F1:CTL vs F1:CCH, p = 0.0138; Figures 3C, 3E). However, in F0 animals directly treated with RHC prior to CCH induction, such recognition memory impairment was prevented, with RHC-treated CCH mice exhibiting a significant increase in DI compared to CCH-alone animals (F0:RHC + CCH vs F0:CCH, p = 0.0151; Figure 3C). Of note, the F1 offspring of RHC-treated animals also exhibited similar resilience to CCH-induced memory loss with DI values significantly increased relative to control offspring derived from untreated parents ([F1:RHC]* + CCH vs F1:CCH, p = 0.0003; Figure 3E). Thus, RHC prevented CCH-induced losses in object recognition memory in animals receiving RHC treatment directly, as well as in their untreated offspring (Figures 3C and 3E).

### Intergenerational preservation of hippocampal long-term potentiation (LTP) via repetitive hypoxic conditioning (RHC)

Application of tetanus led to an initial increase in the slope of field excitatory post-synaptic potential (fEPSP) in all groups (Figures 4B, 4D), with the average of the last twenty minutes post-TBS (40 - 60 min post-TBS) represented in Figures 4B and 4D. Two-way ANOVA revealed a significant main effect of RHC in the F0 generation (F1,34 = 5.071, p = 0.0309) and F1 generation (F_1,22_ = 4.857, p = 0.0383), but no RHC X CCH interaction (Figures 4C, 4E). These data indicate that RHC induces a significant enhancement in LTP compared to untreated animals. Although there was a trend toward a reduction of LTP in CCH animals compared to controls, two-way ANOVA revealed no main effect of CCH. However, a Mann-Whitney test revealed a significant reduction of LTP in CCH animals relative to controls (F0: CTL vs F0: CCH, p = 0.0242; F1: CTL vs F1: CCH, p = 0.0047), suggesting that 4 months of CCH decreased LTP.

### *In vivo* measures of object recognition memory correlate with *ex vivo* measures of synaptic plasticity

In both F0 and F1 animal groups, there was a significant correlation (r = 0.8095, p = 0.0218 for F0; r = 0.8009, p = 0.0009 for F1; Spearman Rank Test) between novel object preference discrimination indices (DI) and averaged long-term potentiation (LTP) field excitatory postsynaptic potential (fEPSP) magnitudes, such that animals that exhibited higher DI also demonstrated heightened synaptic plasticity (Figures 5A, 5B).

**Figure 5.**
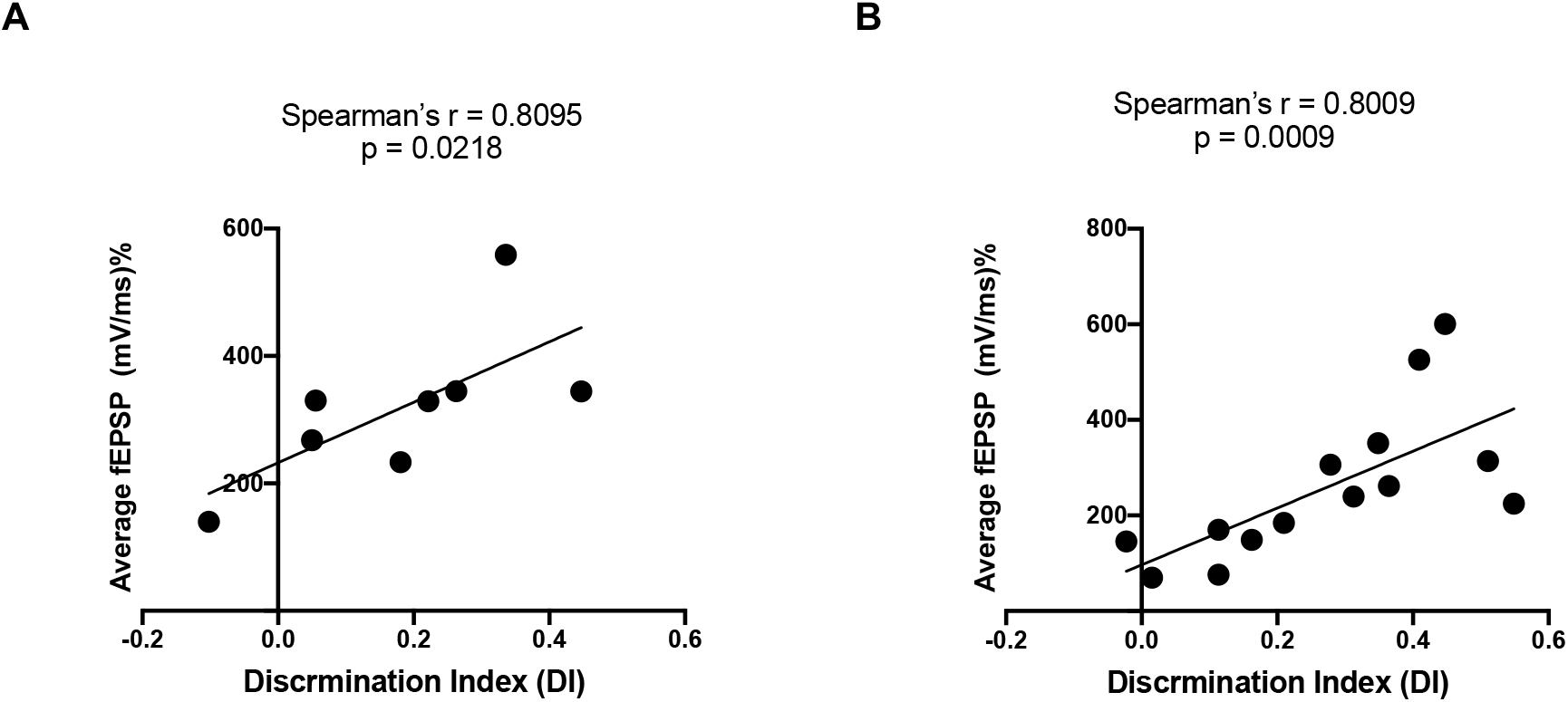
Correlation between *in vivo* measures of object recognition memory and *ex vivo* measurements of synaptic plasticity. There was a significant correlation between discrimination indices (measured *in vivo* by novel object preference) and averaged long-term potentiation (LTP; measured *ex vivo* by fEPSP magnitude) in both F0 (A) and F1 mice (B). Spearman’s correlation coefficients and p-values shown for each generation.

## DISCUSSION

Herein we provide the first evidence for the heritability of an epigenetically-induced resilience to dementia. Specifically, we show in a well-established mouse model of VCID that the protection afforded by an epigenetics-based treatment against two fundamental, functional metrics of memory impairment following 3-4 months of disease is manifested not only in treated animals, but also in their first-generation offspring. Such findings support the concept of adaptive epigenetic conditioning as a therapeutic strategy for reducing the burden of dementia both within and across generations.

Given that impairments in NOP test outcomes are observed after 1 mo of CCH in the mouse model of VCID we used (Patel et al., 2017; Dominguez et al., 2018; Miyanohara et al., 2018; Ben-Ari et al., 2019), it is not surprising that, in our study, 3 mo of CCH resulted in significant decreases in discrimination indices and novelty preferences based on exploration time. Important are our findings that both RHC-treated mice and the adult F1 offspring of RHC-treated males and females were resilient to such CCH-induced deficits, with both cohorts exhibiting no loss of recognition memory. Collectively, these results show that an intermittently presented, nonharmful epigenetic stimulus can induce a long-lasting resilience against object recognition memory impairments in the setting of disease, in animals directly treated with RHC as well as in their F1 offspring.

As with the reductions in novel object preference that we observed following 3 months of CCH, we also expected CCH to adversely affect LTP, a well-established electrophysiologic correlate of synaptic plasticity. The lack of a statistically significant reduction in fEPSP amplitudes after 4 months of CCH suggests that hippocampal injury may not be very advanced at this time, in this particular VCID model. Indeed, Nishio et al. (2010) demonstrated that hippocampal atrophy only becomes evident in carotid-stenosed mice after 5-6 mo of CCH. Moreover, unlike AD patients in which hippocampal atrophy is an early diagnostic indicator, in vascular dementia patients, hippocampal injury occurs at a more intermediate extent and at a later time point (Mueller et al., 2010; Pol et al., 2011). Documenting CCH-induced decrements in synaptic plasticity may also relate to the choice of statistical test we used: A statistically significant reduction in LTP was confirmed in mice with 4 mo of CCH relative to controls when assessed by the nonparametric Mann-Whitney U test, but these between-group differences were not robust enough to reach statistical significance using two-way ANOVA across all four experimental groups. Similarly, two-way ANOVA did not reveal a statistically significant interaction between RHC and CCH with respect to RHC enhancing LTP, but observationally the differences in response amplitudes between untreated and RHC-treated groups is clear. Moreover, within-animal correlations between quantitative, *in vivo* assessments of recognition memory and quantitative, *ex vivo* measures of LTP, were strong in mice from both generations.

Overall, our findings in a chronic neurodegenerative disease model suggest that the efficacy of appropriately titrated conditioning-based therapeutics can extend beyond acute cerebral injuries like stroke and trauma (Stetler et al., 2014; Gidday, 2015; Verges et al., 2015; Baillieul et al., 2017; Sprick et al., 2019; Hao et al., 2020). Indeed, there is evidence that repetitive patterns of hypoxic conditioning lessen disease severity in preclinical models of AD. Specifically, exposing rats to hypobaric hypoxia for 4h/day for 2 wks reduced memory loss induced by amyloid beta injections in a conditioned, passive avoidance reflex test, and also decreased oxidative stress and neuronal degeneration (Manukhina et al., 2016). A number of neurocognitive metrics were improved as a result of treating 9-mo-old APP/PS1 mice with 2 wks of intermittent hypoxia (also 4h/day), which were associated with increases in neurogenesis and BDNF expression (Meng et al., 2020). Finally, in 6-mo-old triple-transgenic AD mice harboring the PS1/M146V, APPswe, and tauP301L transgenes, a 2-wk period of intermittent hypoxic conditioning similar in design to what we employed here prevented learning and memory deficits, and reduced cortical Aβ levels, when measured 2.5-3.0 mo after treatment (Correia et al., 2021).

With respect to VCID, we are aware of only three labs (Ren et al., 2008; Khan et al., 2015, 2018; Li et al., 2017) that have investigated the pre-clinical efficacy of conditioning-based therapeutics, with all of them using ‘remote limb ischemic conditioning’ (RLIC) as a treatment for CCH-induced cognitive impairment. RLIC typically involves the daily induction of 3-4 successive cycles of a ~5 min period of blood flow reduction to limb skeletal musculature, and boasts a long preclinical history for protecting against ischemic injury in brain, heart, and other tissues that are physically “remote” from the intermittently ischemic skeletal muscle (Hess et al., 2015). Treating rats with RLIC for 4 (Li et al., 2017) or 6 wks (Ren et al., 2008) following complete bilateral carotid artery occlusion resulted in improved Morris Water Maze metrics of spatial learning and memory when measured at the end of treatment. In a mouse model of VCID secondary to microcoil bilateral carotid artery stenosis – the same model we used herein – the Hess lab showed that treating young male mice daily with RLIC for two weeks, initiated one week after microcoil-induced CCH, improved cognitive function and reduced white matter pathology measured at 1 mo (Khan et al., 2015). In a follow up study, daily RLIC treatments for either 1 mo or 4 mo provided essentially equivalent protection across several neurocognitive and neurovascular metrics when these outcomes were ultimately assessed after 6 months of CCH (Khan et al., 2018).

In fact, recent clinical trials report that RLIC improved visuospatial and executive ability in patients with cerebral small-vessel disease ([2x/day for 1 year] Wang Y et al., 2017), and improved cognitive function, as assessed by a neuropsychological battery, in patients with subcortical ischemic vascular dementia ([2x/day for 6 mo] Liao et al., 2019). And intermittent hypoxia (3x/week for 8 wks) improved short-term memory and attention in elderly patients with amnestic mild cognitive impairment (Wang H et al., 2020). Despite their small sample sizes, studies like these, and others of patients suffering from chronic cerebral circulation insufficiency (see You et al., 2019 for review), reflect a growing awareness that distinct temporal patterns of changes in blood oxygenation can trigger epigenetic responses in brain, and perhaps the immune system as well, that promote resilience to the progressive dementia suffered by individuals with AD and VCID. The evidence for environmental factors contributing to the genetic and nongenetic risk of AD and related dementias continues to grow (Finch & Kulminski 2019; Bertogliat et al., 2020; Burtscher et al. 2021); despite the aforementioned studies on RLIC, recognition that the “exposome” can also promote disease resilience has yet to gain as strong of a foothold (Lourida et al., 2019; Dhana 2020; Vineis et al., 2020).

That the aforementioned 1 mo of RLIC in mice resulted in cognition-protective effects against CCH that were sustained at least 5 months thereafter (Khan et al., 2018) is consistent with accumulating evidence that epigenetically induced phenotypic change may, under some conditions, be very long-lasting. Ours is the first study to test the possibility that a dementia-resilient phenotype induced by repetitive epigenetic conditioning can become so ‘hard-wired’ that it is actually passed on to one’s progeny. That the adult, first-generation offspring of our RHC-treated mice exhibited neurobehavioral- and LTP-based metrics of intact memory – despite a duration of CCH that impaired these same metrics in their F1 control counterparts born to untreated parents – documents the heritability of dementia resilience acquired secondary to our RHC treatment. While there is a growing literature on such intergenerational inheritance of epigenetically-modified phenotypes (Skvortsova et al., 2018; Cavalli & Heard, 2019; Perez & Lehner, 2019), and companion studies that are beginning to unravel the mechanisms by which germ cell DNA becomes epigenetically modified and passed to the zygote (Fraser & Lin, 2016; Lacal & Ventura, 2018;; Perez & Lehner, 2019; Sharma 2019; Tuscher & Day, 2019), at a conceptual level, the vast majority of research in this field is focused almost exclusively on the inheritance of unhealthy phenotypes, successfully linking repetitively presented stressors to the inheritance of increased disease susceptibility and/or disease itself (Nelson & Nadeau, 2010; Guerrero-Bosagna & Skinner, 2012; Bohacek & Mansuy, 2013; Saab & Mansuy, 2014; Cavelli & Heard, 2019; Perez & Lerner, 2019). To our knowledge, there are only three mammalian studies that have documented the inheritance of beneficial phenotypes in their adult F1 offspring, but these relate to steady-state, resting health metrics, not protection against disease. Specifically, providing F0 parents with novel objects and exercise wheels improved baseline memory in their F1 offspring (Arai et al., 2009; Benito et al., 2018), and exposure of newborn male mice to unpredictable maternal separation and unpredictable maternal stress favored goal-directed behaviors and behavioral flexibility in their adult offspring (Gapp et al., 2014). Again, the present study is the first to document the inheritance of an epigenetically-induced phenotype providing disease resilience.

Our findings provide strong functional evidence, at both an *in vivo* neurobehavioral level and an *ex vivo* molecular level, of the intergenerational inheritance of an epigenetically-induced resilience to dementia. Although we documented that RHC protects against CCH-induced demyelination in the white matter tracts that define the corpus callosum, we did not directly examine any of the multi-generational, multi-tissue, and multi-cellular mechanisms underlying the heritability of this acquired, disease-protective phenotype. Potentially all of the three primary epigenetic regulatory mechanisms – DNA methylation, histone post-translational modifications, and long-noncoding RNAs, are stimulated by RHC and thus contribute to modulating gene expression in both somatic and germ cells of the F0 mice, and the somatic cells of their F1 progeny (Harrison et al., 2016; Zhang et al., 2018). In an earlier genome-wide methylome analysis of the adult mouse brain prefrontal cortex, we found that genes that become differentially methylated in response to RHC are largely distinct from the differentially-methylated genes that define the cortex of their untreated adult progeny. To our surprise, bioinformatic analyses of these methylomes implicated gene expression changes across a much larger network of broad-based, multifunctional cytoprotective genes in the F1 generation, relative to a smaller, ischemia-protective network of genes in the F0 generation, suggesting that intergenerational transfer of an acquired phenotype to offspring does not necessarily require the faithful recapitulation of the conditioning-modified DNA methylome of the parent (Belmonte et al., 2020). Ultimately, uncovering the proteomic, lipidomic, metabolomic, and other cellular- and regional (e.g., prefrontal cortex, hippocampus)- ‘omic features that define the dementia-resilient phenotype will be critical to the identification of specific downstream therapeutic effectors that protect against memory loss; that said, systemic treatments like RHC and RLIC that trigger multiple responses in tissues may be considered a ‘cocktail’ therapeutic that will not likely be mimicked by single molecule-targeted therapeutics. While extended periods of hypoxia can be pathologic, appropriately titrated intermittent hypoxia shows efficacy as a therapeutic for a number of diseases, without evidence of injury (Navarrette-Opazo 2014; Burtscher et al., 2021). In fact, we showed in our methylome study that even 4 months of RHC – twice the duration as used herein – did not adversely affect the viability of CA1 pyramidal cells, known to be the most sensitive neurons in the brain to hypoxia (Belmonte et al., 2020). In the present study, we found no change in white matter myelin density in RHC-treated mice, again suggesting that our epigenetic stimulus was nonharmful, and yet robustly efficacious in triggering gene expression changes sufficient to induce resilience to memory loss in the face of chronic hypoperfusion.

Some limitations of our study are worthy of mention as they relate to potential future studies. Both parents received our RHC treatment prior to mating; whether epigenetic changes in both eggs and sperm are necessary for the heritability of the dementia-resilient phenotype we documented herein will have to be tested. Interestingly, many studies suggest a paternal origin for intergenerational transmission (Rando 2012; Saben et al., 2016). Whether dementia resilience induced by RHC is manifested in the F2 generation is also unknown. We also did not cross-foster the F1 pups to obviate the remote possibility that RHC enhanced parental behaviors, particularly maternal rearing behaviors, in such a way as to contribute to, or wholly account for, the dementia resilience we documented in their untreated adult offspring.

In conclusion, our study is not only the first to document that brief, repetitive exposures to systemic hypoxia can epigenetically reprogram gene expression to establish a dementia-resilient phenotype, but also that germ cells become epigenetically modified by this stimulus as well, resulting in adult offspring that are also protected against memory loss despite sustained cerebral hypoperfusion. The preservation of *in vivo* and *ex vivo* functional metrics of memory/synaptic plasticity we demonstrate herein represents the first mammalian evidence for the inheritance of an induced, disease-resilient, cerebroprotective phenotype. Activating a broad, endogenously-driven resiliency to memory loss – rather than attempting to inhibit specific pathological mechanisms with an exogenous pharmaceutical – is a concept worthy of further investigation, particularly in light of our finding that such an epigenetic therapy may also promote brain health across generations.

## REFERENCES

Ahn, S. M., Kim, Y. R., Kim, H. N., Shin, Y.-I., Shin, H. K., & Choi, B. T. (2016). Electroacupuncture ameliorates memory impairments by enhancing oligodendrocyte regeneration in a mouse model of prolonged cerebral hypoperfusion. Scientific Reports, 6, 28646. https://doi.org/10.1038/srep28646

Antunes, M., & Biala, G. (2012). The novel object recognition memory: Neurobiology, test procedure, and its modifications. Cognitive Processing, 13(2), 93–110. https://doi.org/10.1007/s10339-011-0430-z

Arai, J. A., Li, S., Hartley, D. M., & Feig, L. A. (2009). Transgenerational Rescue of a Genetic Defect in Long-Term Potentiation and Memory Formation by Juvenile Enrichment. The Journal of Neuroscience : The Official Journal of the Society for Neuroscience, 29(5), 1496–1502. https://doi.org/10.1523/JNEUROSCI.5057-08.2009

Baillieul, S., Chacaroun, S., Doutreleau, S., Detante, O., Pépin, J., & Verges, S. (2017). Hypoxic conditioning and the central nervous system: A new therapeutic opportunity for brain and spinal cord injuries? Experimental Biology and Medicine, 242(11), 1198–1206. https://doi.org/10.1177/1535370217712691

Belmonte, K. C. D., Harman, J. C., Lanson, N. A., & Gidday, J. M. (2019). Intra- and intergenerational changes in the cortical DNA methylome in response to therapeutic intermittent hypoxia in mice. Physiological Genomics, 52(1), 20–34. https://doi.org/10.1152/physiolgenomics.00094.2019

Ben-Ari, H., Lifschytz, T., Wolf, G., Rigbi, A., Blumenfeld-Katzir, T., Merzel, T. K., Koroukhov, N., Lotan, A., & Lerer, B. (2019). White matter lesions, cerebral inflammation and cognitive function in a mouse model of cerebral hypoperfusion. Brain Research, 1711, 193–201. https://doi.org/10.1016/j.brainres.2019.01.017

Benito, E., Kerimoglu, C., Ramachandran, B., Pena-Centeno, T., Jain, G., Stilling, R. M., Islam, M. R., Capece, V., Zhou, Q., Edbauer, D., Dean, C., & Fischer, A. (2018). RNA-Dependent Intergenerational Inheritance of Enhanced Synaptic Plasticity after Environmental Enrichment. Cell Reports, 23(2), 546–554. https://doi.org/10.1016/j.celrep.2018.03.059

Bertogliat, M. J., Morris-Blanco, K. C., & Vemuganti, R. (2020). Epigenetic mechanisms of neurodegenerative diseases and acute brain injury. Neurochemistry International, 133, 104642. https://doi.org/10.1016/j.neuint.2019.104642

Bohacek, J., & Mansuy, I. M. (2013). Epigenetic Inheritance of Disease and Disease Risk. Neuropsychopharmacology, 38(1), 220–236. https://doi.org/10.1038/npp.2012.110

Bowers, M. E., & Yehuda, R. (2016). Intergenerational Transmission of Stress in Humans. Neuropsychopharmacology: Official Publication of the American College of Neuropsychopharmacology, 41(1), 232–244. https://doi.org/10.1038/npp.2015.247

Burtscher, J., Maglione, V., Di Pardo, A., Millet, G. P., Schwarzer, C., & Zangrandi, L. (2021). A Rationale for Hypoxic and Chemical Conditioning in Huntington’s Disease. International Journal of Molecular Sciences, 22(2). https://doi.org/10.3390/ijms22020582

Cavalli, G., & Heard, E. (2019). Advances in epigenetics link genetics to the environment and disease. Nature, 571(7766), 489–499. https://doi.org/10.1038/s41586-019-1411-0

Coltman, R., Spain, A., Tsenkina, Y., Fowler, J. H., Smith, J., Scullion, G., Allerhand, M., Scott, F., Kalaria, R. N., Ihara, M., Daumas, S., Deary, I. J., Wood, E., McCulloch, J., & Horsburgh, K. (2011). Selective white matter pathology induces a specific impairment in spatial working memory. Neurobiology of Aging, 32(12), 2324.e7–2324.e12. https://doi.org/10.1016/j.neurobiolaging.2010.09.005

Correia, S. C., Machado, N. J., Alves, M. G., Oliveira, P. F., & Moreira, P. I. (2021). Intermittent Hypoxic Conditioning Rescues Cognition and Mitochondrial Bioenergetic Profile in the Triple Transgenic Mouse Model of Alzheimer’s Disease. International Journal of Molecular Sciences, 22(1), 461. https://doi.org/10.3390/ijms22010461

Dhana, K., Evans, D. A., Rajan, K. B., Bennett, D. A., & Morris, M. C. (2020). Healthy lifestyle and the risk of Alzheimer dementia: Findings from 2 longitudinal studies. Neurology, 95(4), e374–e383. https://doi.org/10.1212/WNL.0000000000009816

Dominguez, R., Zitting, M., Liu, Q., Patel, A., Babadjouni, R., Hodis, D. M., Chow, R. H., & Mack, W. J. (2018). Estradiol Protects White Matter of Male C57BL6J Mice against Experimental Chronic Cerebral Hypoperfusion. Journal of Stroke and Cerebrovascular Diseases, 27(7), 1743–1751. https://doi.org/10.1016/j.jstrokecerebrovasdis.2018.01.030

Finch, C. E., & Kulminski, A. M. (2019). The Alzheimer’s Disease Exposome. Alzheimer’s & Dementia: The Journal of the Alzheimer’s Association, 15(9), 1123–1132. https://doi.org/10.1016/j.jalz.2019.06.3914

Fraser, R., & Lin, C.-J. (2016). Epigenetic reprogramming of the zygote in mice and men: On your marks, get set, go! Reproduction (Cambridge, England), 152(6), R211–R222. https://doi.org/10.1530/REP-16-0376

Gapp, K., Soldado-Magraner, S., Alvarez-Sánchez, M., Bohacek, J., Vernaz, G., Shu, H., Franklin, T. B., Wolfer, D., & Mansuy, I. M. (2014). Early life stress in fathers improves behavioural flexibility in their offspring. Nature Communications, 5, 5466. https://doi.org/10.1038/ncomms6466

Gidday, J. M. (2015). Extending Injury- and Disease-Resistant CNS Phenotypes by Repetitive Epigenetic Conditioning. Frontiers in Neurology, 6. https://doi.org/10.3389/fneur.2015.00042

Gorelick, P. B., Counts, S. E., & Nyenhuis, D. (2016). Vascular cognitive impairment and dementia. Biochimica et Biophysica Acta, 1862(5), 860–868. https://doi.org/10.1016/j.bbadis.2015.12.015

Guerrero-Bosagna, C., & Skinner, M. K. (2012). Environmentally induced epigenetic transgenerational inheritance of phenotype and disease. Molecular and Cellular Endocrinology, 354(1), 3–8. https://doi.org/10.1016/j.mce.2011.10.004

Hao, Y., Xin, M., Feng, L., Wang, X., Wang, X., Ma, D., & Feng, J. (2020). Review Cerebral Ischemic Tolerance and Preconditioning: Methods, Mechanisms, Clinical Applications, and Challenges. Frontiers in Neurology, 11. https://doi.org/10.3389/fneur.2020.00812

Harman, J. C., Guidry, J. J., & Gidday, J. M. (2020). Intermittent Hypoxia Promotes Functional Neuroprotection from Retinal Ischemia in Untreated First-Generation Offspring: Proteomic Mechanistic Insights. Investigative Ophthalmology & Visual Science, 61(11), 15. https://doi.org/10.1167/iovs.61.11.15

Harrison, J. S., Cornett, E. M., Goldfarb, D., DaRosa, P. A., Li, Z. M., Yan, F., Dickson, B. M., Guo, A. H., Cantu, D. V., Kaustov, L., Brown, P. J., Arrowsmith, C. H., Erie, D. A., Major, M. B., Klevit, R. E., Krajewski, K., Kuhlman, B., Strahl, B. D., & Rothbart, S. B. (2016). Hemi-methylated DNA regulates DNA methylation inheritance through allosteric activation of H3 ubiquitylation by UHRF1. ELife, 5, e17101. https://doi.org/10.7554/eLife.17101

Heron, M. (2019). Deaths: Leading Causes for 2017. National Vital Statistics Reports, 68(6), 77.

Hess, D., Khan, M., Morgan, J., & Hoda, M. N. (2015). Remote ischemic conditioning: A treatment for vascular cognitive impairment. Brain Circulation, 1(2), 133. https://doi.org/10.4103/2394-8108.172885

Iadecola, C., Duering, M., Hachinski, V., Joutel, A., Pendlebury, S. T., Schneider, J. A., & Dichgans, M. (2019). Vascular Cognitive Impairment and Dementia: JACC Scientific Expert Panel. Journal of the American College of Cardiology, 73(25), 3326–3344. https://doi.org/10.1016/j.jacc.2019.04.034

Kalaria, R. N., Akinyemi, R., & Ihara, M. (2016). Stroke injury, cognitive impairment and vascular dementia. Biochimica et Biophysica Acta, 1862(5), 915–925. https://doi.org/10.1016/j.bbadis.2016.01.015

Khan, M. B., Hafez, S., Hoda, Md. N., Baban, B., Wagner, J., Awad, M. E., Sangabathula, H., Haigh, S., Elsalanty, M., Waller, J. L., & Hess, D. C. (2018). Chronic Remote Ischemic Conditioning Is Cerebroprotective and Induces Vascular Remodeling in a VCID Model. Translational Stroke Research, 9(1), 51–63. https://doi.org/10.1007/s12975-017-0555-1

Khan, M. B., Hoda, M. N., Vaibhav, K., Giri, S., Wang, P., Waller, J. L., Ergul, A., Dhandapani, K. M., Fagan, S. C., & Hess, D. C. (2015). Remote Ischemic Postconditioning: Harnessing Endogenous Protection in a Murine Model of Vascular Cognitive Impairment. Translational Stroke Research, 6, 69–77. https://doi.org/10.1007/s12975-014-0374-6

Lacal, I., & Ventura, R. (2018). Epigenetic Inheritance: Concepts, Mechanisms and Perspectives. Frontiers in Molecular Neuroscience, 11. https://doi.org/10.3389/fnmol.2018.00292

Li, X., Ren, C., Li, S., Han, R., Gao, J., Huang, Q., Jin, K., Luo, Y., & Ji, X. (2017). Limb Remote Ischemic Conditioning Promotes Myelination by Upregulating PTEN/Akt/mTOR Signaling Activities after Chronic Cerebral Hypoperfusion. Aging and Disease, 8(4), 392–401. https://doi.org/10.14336/AD.2016.1227

Liao, Z., Bu, Y., Li, M., Han, R., Zhang, N., Hao, J., & Jiang, W. (2019). Remote ischemic conditioning improves cognition in patients with subcortical ischemic vascular dementia. BMC Neurology, 19(1), 206. https://doi.org/10.1186/s12883-019-1435-y

Lourida, I., Hannon, E., Littlejohns, T. J., Langa, K. M., Hyppönen, E., Kuzma, E., & Llewellyn, D. J. (2019). Association of Lifestyle and Genetic Risk With Incidence of Dementia. JAMA, 322(5), 430–437. https://doi.org/10.1001/jama.2019.9879

Lueptow, L. M. (2017). Novel Object Recognition Test for the Investigation of Learning and Memory in Mice. Journal of Visualized Experiments : JoVE, 126. https://doi.org/10.3791/55718

Manukhina, E. B., Downey, H. F., Shi, X., & Mallet, R. T. (2016). Intermittent hypoxia training protects cerebrovascular function in Alzheimer’s disease. Experimental Biology and Medicine, 241(12), 1351–1363. https://doi.org/10.1177/1535370216649060

Meng, S.-X., Wang, B., & Li, W.-T. (2020). Intermittent hypoxia improves cognition and reduces anxiety-related behavior in APP/PS1 mice. Brain and Behavior, 10(2), e01513. https://doi.org/10.1002/brb3.1513

Miki, K., Ishibashi, S., Sun, L., Xu, H., Ohashi, W., Kuroiwa, T., & Mizusawa, H. (2009). Intensity of chronic cerebral hypoperfusion determines white/gray matter injury and cognitive/motor dysfunction in mice. Journal of Neuroscience Research, 87(5), 1270–1281. https://doi.org/10.1002/jnr.21925

Miller, B. A., Perez, R. S., Shah, A. R., Gonzales, E. R., Park, T. S., & Gidday, J. M. (2001). Cerebral protection by hypoxic preconditioning in a murine model of focal ischemia-reperfusion. Neuroreport, 12(8), 1663–1669.

Miyanohara, J., Kakae, M., Nagayasu, K., Nakagawa, T., Mori, Y., Arai, K., Shirakawa, H., & Kaneko, S. (2018). TRPM2 Channel Aggravates CNS Inflammation and Cognitive Impairment via Activation of Microglia in Chronic Cerebral Hypoperfusion. The Journal of Neuroscience, 38(14), 3520–3533. https://doi.org/10.1523/JNEUROSCI.2451-17.2018

Moreno-Castilla, P., Guzman-Ramos, K., & Bermudez-Rattoni, F. (2018). Chapter 28—Object Recognition and Object Location Recognition Memory – The Role of Dopamine and Noradrenaline. In A. Ennaceur & M. A. de Souza Silva (Eds.), Handbook of Behavioral Neuroscience (Vol. 27, pp. 403–413). Elsevier. https://doi.org/10.1016/B978-0-12-812012-5.00028-8

Mueller, S. G., Schuff, N., Yaffe, K., Madison, C., Miller, B., & Weiner, M. W. (2010). Hippocampal atrophy patterns in mild cognitive impairment and Alzheimer’s disease. Human Brain Mapping, 31(9), 1339–1347. https://doi.org/10.1002/hbm.20934

Navarrete-Opazo, A., & Mitchell, G. S. (2014). Therapeutic potential of intermittent hypoxia: A matter of dose. American Journal of Physiology-Regulatory, Integrative and Comparative Physiology, 307(10), R1181–R1197. https://doi.org/10.1152/ajpregu.00208.2014

Nelson, V. R., & Nadeau, J. H. (2010). Transgenerational genetic effects. Epigenomics, 2(6), 797–806. https://doi.org/10.2217/epi.10.57

Nicoll, R. A. (2017). A Brief History of Long-Term Potentiation. Neuron, 93(2), 281–290. https://doi.org/10.1016/j.neuron.2016.12.015

Nishio, K., Ihara, M., Yamasaki, N., Kalaria, R. N., Maki, T., Fujita, Y., Ito, H., Oishi, N., Fukuyama, H., Miyakawa, T., Takahashi, R., & Tomimoto, H. (2010). A Mouse Model Characterizing Features of Vascular Dementia With Hippocampal Atrophy. Stroke, 41(6), 1278–1284. https://doi.org/10.1161/STROKEAHA.110.581686

O’Brien, J. T., & Thomas, A. (2015). Vascular dementia. The Lancet, 386(10004), 1698–1706. https://doi.org/10.1016/S0140-6736(15)00463-8

Orso, R., Creutzberg, K. C., Wearick-Silva, L. E., Wendt Viola, T., Tractenberg, S. G., Benetti, F., & Grassi-Oliveira, R. (2019). How Early Life Stress Impact Maternal Care: A Systematic Review of Rodent Studies. Frontiers in Behavioral Neuroscience, 13. https://doi.org/10.3389/fnbeh.2019.00197

Patel, A., Moalem, A., Cheng, H., Babadjouni, R. M., Patel, K., Hodis, D. M., Chandegara, D., Cen, S., He, S., Liu, Q., & Mack, W. J. (2017). Chronic cerebral hypoperfusion induced by bilateral carotid artery stenosis causes selective recognition impairment in adult mice. Neurological Research, 39(10), 910–917. https://doi.org/10.1080/01616412.2017.1355423

Pembrey, M., Saffery, R., Bygren, L. O., Network in Epigenetic Epidemiology, & Network in Epigenetic Epidemiology. (2014). Human transgenerational responses to early-life experience: Potential impact on development, health and biomedical research. Journal of Medical Genetics, 51(9), 563–572. https://doi.org/10.1136/jmedgenet-2014-102577

Perez, M. F., & Lehner, B. (2019). Intergenerational and transgenerational epigenetic inheritance in animals. Nature Cell Biology, 21(2), 143–151. https://doi.org/10.1038/s41556-018-0242-9

Pol, L. van de, Gertz, H.-J., Scheltens, P., & Wolf, H. (2011). Hippocampal Atrophy in Subcortical Vascular Dementia. Neurodegenerative Diseases, 8(6), 465–469. https://doi.org/10.1159/000326695

Rando, O. J. (2012). Daddy Issues: Paternal Effects on Phenotype. Cell, 151(4), 702–708. https://doi.org/10.1016/j.cell.2012.10.020

Ren, C., Gao, X., Steinberg, G. K., & Zhao, H. (2008). Limb remote-preconditioning protects against focal ischemia in rats and contradicts the dogma of therapeutic time windows for preconditioning. Neuroscience, 151(4), 1099–1103. https://doi.org/10.1016/j.neuroscience.2007.11.056

Saab, B. J., & Mansuy, I. M. (2014). Neurobiological disease etiology and inheritance: An epigenetic perspective. Journal of Experimental Biology, 217(1), 94–101. https://doi.org/10.1242/jeb.089995

Saben, J. L., Boudoures, A. L., Asghar, Z., Thompson, A., Drury, A., Zhang, W., Chi, M., Cusumano, A., Scheaffer, S., & Moley, K. H. (2016). Maternal Metabolic Syndrome Programs Mitochondrial Dysfunction via Germline Changes across Three Generations. Cell Reports, 16(1), 1–8. https://doi.org/10.1016/j.celrep.2016.05.065

Sharma, U. (2019). Paternal Contributions to Offspring Health: Role of Sperm Small RNAs in Intergenerational Transmission of Epigenetic Information. Frontiers in Cell and Developmental Biology, 7. https://doi.org/10.3389/fcell.2019.00215

Shibata, M., Ohtani, R., Ihara, M., & Tomimoto, H. (2004). White matter lesions and glial activation in a novel mouse model of chronic cerebral hypoperfusion. Stroke, 35(11), 2598–2603. https://doi.org/10.1161/01.STR.0000143725.19053.60

Shibata, M., Yamasaki, N., Miyakawa, T., Kalaria, R. N., Fujita, Y., Ohtani, R., Ihara, M., Takahashi, R., & Tomimoto, H. (2007). Selective impairment of working memory in a mouse model of chronic cerebral hypoperfusion. Stroke, 38(10), 2826–2832. https://doi.org/10.1161/STROKEAHA.107.490151

Skvortsova, K., Iovino, N., & Bogdanović, O. (2018). Functions and mechanisms of epigenetic inheritance in animals. Nature Reviews Molecular Cell Biology, 19(12), 774–790. https://doi.org/10.1038/s41580-018-0074-2

Sprick, J. D., Mallet, R. T., Przyklenk, K., & Rickards, C. A. (2019). Ischemic and Hypoxic Conditioning: Potential for Protection of Vital Organs. Experimental Physiology, 104(3), 278–294. https://doi.org/10.1113/EP087122

Stetler, R. A., Leak, R. K., Gan, Y., Li, P., Zhang, F., Hu, X., Jing, Z., Chen, J., Zigmond, M. J., & Gao, Y. (2014). Preconditioning provides neuroprotection in models of CNS disease: Paradigms and clinical significance. Progress in Neurobiology, 114, 58–83. https://doi.org/10.1016/j.pneurobio.2013.11.005

Stowe, A. M., Altay, T., Freie, A. B., & Gidday, J. M. (2011). Repetitive hypoxia extends endogenous neurovascular protection for stroke. Annals of Neurology, 69(6), 975–985. https://doi.org/10.1002/ana.22367

Stowe, A. M., Wacker, B. K., Cravens, P. D., Perfater, J. L., Li, M. K., Hu, R., Freie, A. B., Stüve, O., & Gidday, J. M. (2012). CCL2 upregulation triggers hypoxic preconditioning-induced protection from stroke. Journal of Neuroinflammation, 9, 33. https://doi.org/10.1186/1742-2094-9-33

Sumner, R. L., Spriggs, M. J., Muthukumaraswamy, S. D., & Kirk, I. J. (2020). The role of Hebbian learning in human perception: A methodological and theoretical review of the human Visual Long-Term Potentiation paradigm. Neuroscience & Biobehavioral Reviews, 115, 220–237. https://doi.org/10.1016/j.neubiorev.2020.03.013

Tuscher, J. J., & Day, J. J. (2019). Multigenerational epigenetic inheritance: One step forward, two generations back. Neurobiology of Disease, 132, 104591. https://doi.org/10.1016/j.nbd.2019.104591

Verges, S., Chacaroun, S., Godin-Ribuot, D., & Baillieul, S. (2015). Hypoxic Conditioning as a New Therapeutic Modality. Frontiers in Pediatrics, 3. https://doi.org/10.3389/fped.2015.00058

Vineis, P., Robinson, O., Chadeau-Hyam, M., Dehghan, A., Mudway, I., & Dagnino, S. (2020). What is new in the exposome? Environment International, 143, 105887. https://doi.org/10.1016/j.envint.2020.105887

Vogel-Ciernia, A., & Wood, M. A. (2014). Examining Object Location and Object Recognition Memory in Mice. Current Protocols in Neuroscience, 69(1), 8.31.1–8.31.17. https://doi.org/10.1002/0471142301.ns0831s69

Wang, H., Shi, X., Schenck, H., Hall, J. R., Ross, S. E., Kline, G. P., Chen, S., Mallet, R. T., & Chen, P. (2020). Intermittent Hypoxia Training for Treating Mild Cognitive Impairment: A Pilot Study. American Journal of Alzheimer’s Disease & Other Dementiasr, 35, 1533317519896725. https://doi.org/10.1177/1533317519896725

Wang, Y., Meng, R., Song, H., Liu, G., Hua, Y., Cui, D., Zheng, L., Feng, W., Liebeskind, D. S., Fisher, M., & Ji, X. (2017). Remote Ischemic Conditioning May Improve Outcomes of Patients With Cerebral Small-Vessel Disease. Stroke, 48(11), 3064–3072. https://doi.org/10.1161/STROKEAHA.117.017691

You, J., Feng, L., Bao, L., Xin, M., Ma, D., & Feng, J. (2019). Potential Applications of Remote Limb Ischemic Conditioning for Chronic Cerebral Circulation Insufficiency. Frontiers in Neurology, 10. https://doi.org/10.3389/fneur.2019.00467

Zhang, W., Song, M., Qu, J., & Liu, G.-H. (2018). Epigenetic Modifications in Cardiovascular Aging and Diseases. Circulation Research, 123(7), 773–786. https://doi.org/10.1161/CIRCRESAHA.118.312497

